# Association of variants in *ACE, ACTN3, AGT, IL6* and *BDKRB2* genes with athlete status and playing position in Colombian amateur rugby athletes

**DOI:** 10.1101/2022.05.05.490786

**Authors:** Efraín Paz Garcia, Gerardo David Gonzalez, Guillermo Barreto

## Abstract

Genetic polymorphisms are involved in different metabolic pathways that are manifested at the physiological level and have been associated with specific phenotypes in sport from anthropometric and functional characterizations that pose conditional and physiological demands for the rugby athlete. The identification of this type of polymorphisms in athletes represents a resource that contributes significantly to the processes of training, selection and sports orientation.

The purpose of this study was to describe type and frequencies of allelic and genotypic variants in *ACTN3, ACE, AGT, BDRKB2* and *IL6* genes in sub elite rugby athletes in Colombia. Additionally, the polymorphisms found were compared with a control population, as well as contrasted according to playing position backs and forwards.

In this research, 47 individuals from the Vallecaucana rugby league and 67 from a control group (non-athletes) were sampled. All were analyzed for polymorphisms in the *ACE, AGT, ACTN3, IL6* and *BDKRB2* genes, using the PCR RFLPs technique. The significance of the differences between the experimental and control groups was tested by the *X2* test (p <0.05).

In rugby athletes we found a higher frequency of allele D (0.883) *ACE* gene, allele R (0.63) *ACTN3* gene, allele G (0.819) *IL6* gene, all associated with strength and power sports. There are significant genotypic differences between athletes and the control population in all the genes analyzed and significant allelic differences in the *ACE, ACTN3, BDRKB2* and *IL6* genes. When comparing the playing positions (backs vs. forwards), significant genotypic differences were observed in the *ACTN3, BDRKB2, AGT* and *IL6* genes. At the allelic level, the R and X alleles of the *ACTN3* gene and the I allele of the *ACE* gene show significant differences.

In conclusion, in the polymorphisms analyzed, an association with strength sports, explosive strength and rugby is observed. Significant genotypic and allelic differences were also recorded between the backs and forwards positions, as well as significant differences in the allelic and genotypic structure between the group of athletes and the control population.

## Introduction

Rugby is a field game that demands mobility, agility, strength and muscle power. These vary with positional role and also with competitive level. It is a team, contact and ball sport, of intermittent development and acyclic structure, with essential characteristics of high aerobic fitness, speed, strength, muscle power and agility. (Suárez-Moreno Arrones, & Núñez 2011).

In rugby union players are classified into two main groups: forwards (forwards) and backs (¾ line), each group with different anthropometric and conditional characteristics (Brazier et al., 2018). It is a game where high intensity actions alternate with short recovery periods, involving conditional capacities, such as speed, aerobic power and muscle power, however, it is likely that the ability of players to repeat high intensity efforts intermittently throughout the match, is one of the factors that most conditions the performance of the players, (Lopes et al., 2011).

Thus, in rugby, due to its acyclic nature and intensity, the physical demands are very complex, requiring athletes to have high speed, agility, strength, muscle power and endurance. This added to its great popularity and insertion in elite sports has generated a great investigative interest in terms of improving both training processes, such as sports selection interdisciplinary intervening in them and which could not escape the advances in genetics associated with physical performance and sport, being in recent years the subject of several studies framed in the RugbyGen project (Molecular Genetic Characteristics of Elite Rugby Athletes) led by Shane M Heffernan, Alun G Williams and other collaborators who have characterized anthropometrically and physiologically rugby, besides associating it to polymorphisms such as I/D of the *ACE* gene (RS4646994), R/X of *ACTN3* (RS1815739) they found significant relationships with maximal strength, speed, power and aerobic capacity, in addition to identifying genotypic differences between forwards and backs (Brazier et al., 2018, Heffernan et al., 2015, Heffernan et al., 2016., Bell et al., 2012). Similarly gene polymorphisms such as *FTO* T/N were associated with body composition (Heffernan et al., 2017), *COL5A1* RS12722 C/T and RS3196378 C/A with injury risk (Heffernan et al., 2017), (Brazier et al., 2019), *PPARA* G/C with regulation of lipid and glucose metabolism, associated with power sports and with aerobic and anaerobic system in team sports (Ahmetov et al., 2013, Massidda, Calò, et al.,2019). It was also positively associated the D allele of the *ACE* gene with leg power for forwards and backs finding significant differences, obtaining the latter more relative strength and speed for both ID and DD genotypes (Bell et al., 2010); *ACTN3* R/X was also associated with body composition and percentage of fat and muscle power according to playing position without finding significant differences in these variables (Bell et al., 2012).

This research was proposed to study other genetic variants associated with strength - power sports, as well as genes involved in the same metabolic pathway, is the case of *ACE* (RS4646994), *AGT* (RS699) and *BDKRB2* (RS5810761) genes, associated within the renin, angiotensin, aldosterone and kalikrein kinin pathway, which influence the responses of vasoconstriction and vasodilatation, against physical activity. These two pathways interact thanks to the influence of angiotensin converting enzyme, which transforms angiotensinogen I from angiotensinogen (*AGT* gene) into angiotensinogen II, which is a trigger of vasoconstriction in blood vessels. In addition *ACE* is related to the kalikrein kinin pathway, as it intervenes in the degradation of kinins, also favoring vasoconstriction, by degrading kinins that influence the vasodilator activity of blood vessels, likewise the relationship of modulation of *BDRKB2* on the activity of *ACE* in serum (Alves et al., 2013). It reaffirms the association already made with speed and power athletes both in rugby and other sports of the *ACTN3* R/X gene (Andrade-Mayorga et al., 2019, Pimenta et al., 2012). In another way, the G/C polymorphism of the *IL6* gene (RS1800795) associated with studies with power athletes and creatine kinase values as an indicator of muscle damage that linked the *IL6* GG genotype with great muscle recovery capacity, in addition to referencing it as a genotype that predisposes to explosive strength (Sarzynski et al., 2016, Szalata et al., 2019, Yamin et al., 2008) is associated.

The aim of this study was to identify the allelic and genotypic distribution of *ACE, ACTN3, BDKRB2, AGT* and *IL6* genes in players of the Vallecaucana Rugby League, and to associate this distribution with the athlete status and the positions played in the field of play and its differentiation with a control group.

## Methods

### Participants

A total of 116 individuals were sampled: 47 rugby league players from Valle del Cauca and 69 controls. The criteria to consider an athlete were: belonging to the Rugby Valle-Pro high performance program, and participating in national and international competitions. The participants of the control group were people without experience in competition and recreational athletes with less than three days of training per week. This project was approved by the Human Ethics Committee of the Universidad del Valle, and was carried out in accordance with the declaration of Helsinki, following the protocols required in human studies, according to resolution 008430/1993 of the Colombian Ministry of Health. Before taking the sample, each athlete signed an informed consent form, where he/she agreed to participate in the present study.

### Genotyping

The genetic material was extracted from DNA of buccal epithelial cells, following the protocol outlined by Quinque et al., (2006). Primers amplifying the genes, *ACE* (Cieszczyk. et al., 2010), *ACTN3* (Cięszczyk 2011), *AGT* (Gómez-Gallego 2009), *IL6* (Yamin et al. 2008) and *BDKRB2* (Grenda A. 2014) were used for molecular analyses. PCR reactions were performed in a VeritiTM thermal cycler (Applied Biosystem), at a volume of 25μl, which contained, 3 mM MgCl2, 0.1 mM primers, 0.1 mM dNTPs, 1X buffer, 1X Taq polymerase. The amplified fragments of the AGT and *ACTN3* genes were subjected to the RFLP technique, using the restriction enzymes SfnaI and DdeI exposing the amplified fragments for 12 hours to the enzymatic action. Electrophoresis was performed in 8% polyacrylamide gels in TBE1X, of the products obtained in PCR and enzymatic digestions. These electrophoreses were run at 130 volts for 1 hour and at 250 volts for 2 hours and visualized on gels stained with silver nitrate.

## DATA ANALYSIS

Using the data obtained in the previously described molecular techniques, a database of presence (1) or absence (0) of the polymorphisms found for the *ACE, ACTN3, AGT, IL6* and *BDKRB2* genes was constructed for each individual sampled. With this database, a table of absolute frequencies was created, with which genotypic and allelic frequencies were calculated and placed in their corresponding tables (Elston. et al 2012). The genotypic frequencies of the elite athlete population and the control population were evaluated for compatibility with Hardy-Weinberg equilibrium (Templeton A. 2006). Genotypic distribution and allele frequencies were compared between the elite athlete population and the control population, and genotypic distribution and allele frequencies were compared between athletes and the positions on the field of play they played, associating the positions into two groups, forwards and backs (Weir B. 1996). Significance was tested with the *X2* test (p <0.05), using EXCEL software.

## Results

A predisposition to genotypes related to strength and power was observed, as shown in Table 1, which have been associated with the genotypes DD of the *ACE* gene, RR and RX of the *ACTN3* gene, CC and CT of the *AGT* gene and GG of the *IL6* gene. The *BDKRB2* gene presents a majority of individuals with the −9+9 genotype which is associated with a predisposition to vasodilation linked to aerobic capacity, reported by (Williams et al. 2004) and (Alves et al. 2013) in the reviews of Rankinen et al. (2006) and Szalata et al. (2019).

**Table 1.** Absolute genotypic frequencies of the two populations: Rugby athletes and control for the five genes studied.

|  | ACE |  |  | ACTN3 |  |  | BDKRB2 |  |  | AGT |  |  | IL6 |  |  |
| --- | --- | --- | --- | --- | --- | --- | --- | --- | --- | --- | --- | --- | --- | --- | --- |
|  | DD | II | ID | RR | XX | RX | -9/-9 | +9/+9 | -9/+9 | CC | TT | CT | GG | CC | GC |
| Rugby | 37 | 1 | 9 | 16 | 4 | 27 | 7 | 5 | 35 | 2 | 0 | 45 | 35 | 5 | 7 |
| Controls | 21 | 26 | 22 | 13 | 18 | 38 | 27 | 6 | 36 | 22 | 13 | 34 | 61 | 5 | 3 |

**Graph No 1.**
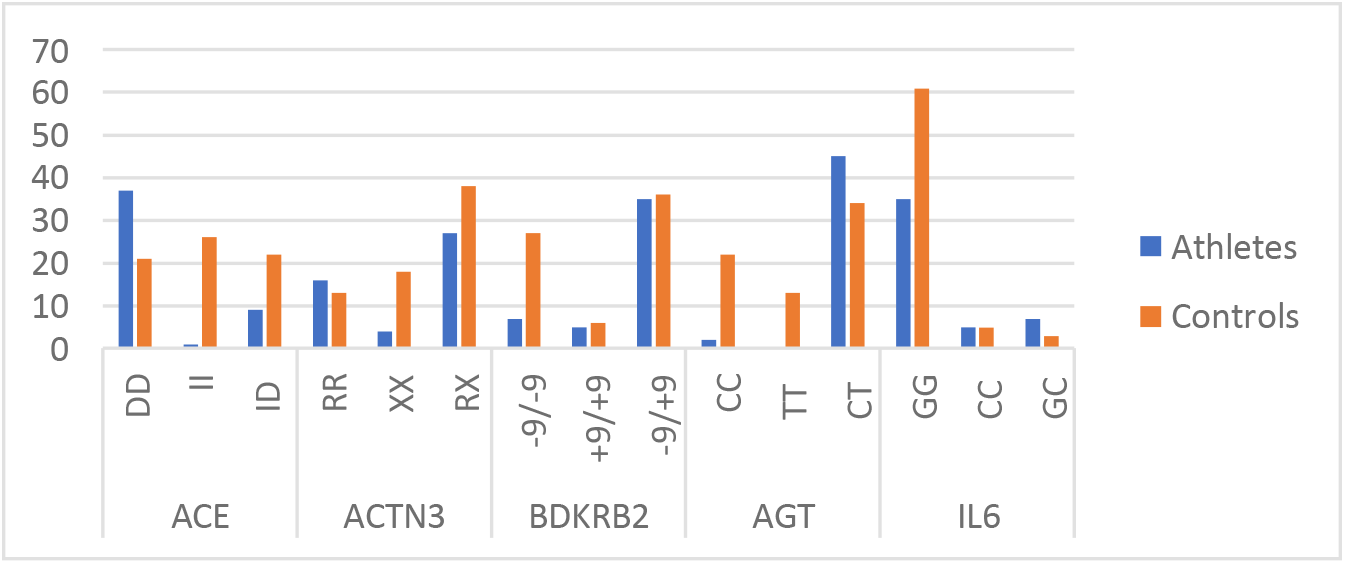
Absolute genotypic frequencies of rugby players for the five genes studied. The abscissa indicates the genotypes per locus and the ordinates the absolute frequency.

Table No. 2 shows the allele frequencies obtained, the ACE gene, presents in the total population of rugby athletes, a majority of the D allele (0.883), which has been reported as a promoter of vasoconstriction and related to strength and power by (Eider et al., 2013) in Polish athletes (Drozdovska et al., 2013) and elite Ukrainian athletes due to increased production of ACE, compared to the I allele. The ACTN3 gene showed a higher frequency of the R allele (0.63), in the athlete population, as well as in the AGT genes the C allele (0.52), BDKRB2 allele −9 (0.521), the interleukin 6 gene, showed a higher presence of the G allele (0.819) in athletes, which has been associated in previous works to power genotypes (Ruiz et al., 2010) and also to a better adaptation to muscle recovery processes (Yamin et al., 2008). In the control population, a greater presence of alleles −9 (0.652) of the BDRKB2 gene, C (0.57) of the AGT gene and G (0.906) of the IL6 gene is observed.

**Table No. 2.** Relative allele frequencies for the populations: rugby players and controls

|  | ACE |  | ACTN3 |  | BDKRB2 |  | AGT |  | IL6 |  |
| --- | --- | --- | --- | --- | --- | --- | --- | --- | --- | --- |
|  | D | I | R | X | -9 | +9 | C | T | G | C |
| **Rugby** | 0,883 | 0,12 | 0,63 | 0,37 | 0,521 | 0,479 | 0,52 | 0,479 | 0,819 | 0,1809 |
| **Controls** | 0,464 | 0,54 | 0,46 | 0,54 | 0,652 | 0,348 | 0,57 | 0,435 | 0,906 | 0,0942 |

The rugby athletes group, presented HW imbalance in IL6, BDKRB2 and AGT genes, while the control group presented HW imbalance for IL6 and ACE genes. (Table No. 3).

**Table No. 3.** Significance of the chi-square test to test the hypothesis of existence of Hardy-Weinberg equilibrium (HWE) for the five genes considering the Rugby athletes population, the control group and the population as a whole (rugby athletes+control group).

|  | HWE P - VALUE (0.05) |  |  |  |  |
| --- | --- | --- | --- | --- | --- |
|  | ACE | ACTN3 | BDKRB2 | AGT | IL6 |
| **Rugby** | 0,6149 | 0,1163 | 0,0007 | 0,0000 | 0,0014 |
| **Controls** | 0,0029 | 0,3729 | 0,2128 | 0,9830 | 0,0002 |

Table 4 shows how there are significant differences when comparing the group of sampled athletes against the control population, at genotypic level all genes show significant differences, at allelic level significant differences are observed in the D and I alleles of the ACE gene, R and X of the ACTN3 gene, +9 allele of the BDRKB2 gene and the C allele of the IL6 gene.

**Table 4.** Comparison of genotypic and allelic frequencies between rugby athletes and the control group.

|  | P ALLELS |  |  |  |  |  |  |  |  |  |
| --- | --- | --- | --- | --- | --- | --- | --- | --- | --- | --- |
|  | ACE<br>D | ACE<br>I | ACTN3<br>R | ACTN3<br>X | BDKRB2<br>-9 | BDKRB2<br>+9 | AGT<br>C | AGT<br>T | AGT<br>G | AGT<br>C |
| **Rugby vs controls** | 0,0000 | 0,0000 | 0,0240 | 0,0358 | 0,1316 | 0,0390 | 0,6184 | 0,5701 | 0,4074 | 0,0102 |
|  | P GENOTYPE |  |  |  |  |  |  |  |  |  |
|  | ACE | ACTN3 | BDKRB2 | AGT | IL6 |  |  |  |  |  |
| **Rugby vs controls** | 0,0000 | 0,0023 | 0,0001 | 0,0000 | 0,0000 |  |  |  |  |  |

When comparing by the *X2* test, the game positions (backs vs forward), it is observed in table 5, significant differences at genotypic level in the genes *ACTN3, BDRKB2, AGT* and *IL6*. At allelic level, only the R and X alleles of the *ACTN3* gene present significant differences and the I allele of the ACE gene, the rest of the genes do not show significant differences at the allelic level, when comparing the gametic positions.

**Table 5.** Comparison of genotypic and allele frequencies between Backs and Forwards positions.

|  | P ALLELS |  |  |  |  |  |  |  |  |  |
| --- | --- | --- | --- | --- | --- | --- | --- | --- | --- | --- |
|  | ACE<br>D | ACE<br>I | ACTN3<br>R | ACTN3<br>X | BDKRB2<br>-9 | BDKRB2<br>+9 | AGT<br>C | AGT<br>T | AGT<br>G | AGT<br>C |
| Backs vs forwards | 0,5859 | 0,0495 | 0,0000 | 0,0013 | 0,9600 | 0,9580 | 0,7686 | 0,7686 | 0,6455 | 0,2598 |
|  | P GENOTYPE |  |  |  |  |  |  |  |  |  |
|  | ACE | ACTN3 | BDKRB2 | AGT | IL6 |  |  |  |  |  |
| Backs vs forwards | 0,1057 | 0,0000 | 0,0042 | 0,0000 | 0,0064 |  |  |  |  |  |

## DISCUSSION

This article is the first in Colombia that associates genetic polymorphisms related to physical performance in rugby players of amateur leagues whose results are in agreement with investigations of the Rugby Gen project whose data agree in the *ACE* (I/D) and *ACTN3* (R/X) genes, as well as exploring new polymorphisms associated with physical performance in strength, power, aerobic capacity and muscle recovery, information that currently affects the morphological, functional and motor characterization of rugby and training processes. The H.W. analysis shows imbalances in controls (ACE and IL6) and athletes (IL6, AGT and BDRKB2) possibly due to the size of the sample and the migrant and multiethnic origin of the population in this area of the country. The IL6 gene, which showed an imbalance in athletes and controls, was more marked.

Regarding the athlete status we found that the athletes presented genotypic distributions associated with strength and power, the predominant genotypes were *ACE* (DD: 0.78), ACTN3 (RR: 0.34 and RX: 0.57), *AGT* (CT: 0.95) and *IL6* (GG: 0.74). At the allelic level, the frequencies of alleles D (0.883) of the *ACE* gene, R (0.63) of *ACTN3*, C (0.52) of *AGT* and G (0.819) of *IL6* were predominant. These results are in agreement with those reported by Bouchard, et al 2004 and Rankinen, et al 2006, 2010 and 2016 who determined association of those genotypes with athlete status. Also studies such as Eider et al., 2013 and Santoro et al., 2019 that related, the first power athletes with a predominance of 87% of the D allele and the second with Brazilian football athletes whose predominance of the D allele was 68%; in addition to numerous studies on the *ACTN3* gene (RX) such as those of Pimenta et al., 2012 and Heffernan et al., 2016 that correspond to the genotypic and allelic findings of this research in terms of association with reactive characteristics. Muscular strength is required by forwards in all aspects of fixed scrum formation, where strength is applied isometrically in the first instance, coordinated later in a team push, backs require speed, agility and power, it is also required in rucks and mauls (play actions), in taking the ball away from the opponent, and is necessary for all players to tackle and shed tackles (Reilly, 1997). In a rugby match there are actions where different manifestations of strength are exerted, so we can positively associate these allelic and genotypic distributions presented, which support the characterization of rugby as a sport of strength and speed.

A significant difference is observed between the athletes sampled against the control group in all genotypic frequencies of the genes studied, as well as in the frequencies of the alleles of the *ACE* (D/I), *ACTN3* (R/X), *BDRKB2* (+9) and *IL6*(C) genes, being related to an artificial selection made in the rugby players, oriented to genotypes that have better performances for the practice of this sport, which is in the alactic anaerobic and lactic anaerobic threshold, unlike the control group which is oriented more to aerobic genotypes.

Regarding the aerobic component, it could be observed, as alleles I of the *ACE* gene and X of the *ACTN3* gene, which have been related to a better performance in aerobic activities (Aksenov & Ilyin, 2017) and (Massidda, Voisin, et al., 2019), who reported in contrast to the D alleles of the *ACE* gene and R of *ACTN3*, that the *ACE* I and *ACTN3* X alleles predominated more in association with cyclic endurance sports, in addition to presenting predisposition to muscle damage in the case of the *ACTN3* X allele (Massidda, Voisin, et al., 2019). Thus, significant differences were presented between the athletes and the control group, which represents in athletes a better performance in strength and power actions, but with a base of the aerobic component, because although, the top priority should be speed. The game is based on strength, power, flexibility and the practice of maximum speed technique. The second capacity is strength endurance. This ability allows the player to work repeatedly at maximum intensities. To be able to do this, the player requires the ability to tolerate high levels of lactic acid and to have muscles with a high capacity to plug and fight acidosis at the muscular level. The third is aerobic endurance. A good ability that allows the body to withstand the fatiguing effects of strength endurance activities, and also allows the player to make repeated speed changes (Casajus 2013). In this sense it was estimated as the −9 allele of the *BDRKB2* gene, is present with greater frequency, referencing vasodilator capacity associated with the fundamental aerobic capacity in the performance of a rugby player (Rett, 1990), (Williams et al., 2004), this coupled with alleles I of the *ACE* gene and X of the *ACTN3* gene, could provide a better aerobic performance than the population average.

When comparing within the group of athletes, if there are differences in the genotypes of the players taking into account their position on the field (Backs and Forwards), a significant difference is observed in the genotypes of backs and forwards in *ACTN3* genes (p=0.0000), presenting a greater presence of RR and RX genotypes in backs than in forwards, this does not occur in studies in rugby as no significant differences are presented, as in the case of Bell et al, 2012 which reports that backs had a slightly higher proportion of the RR genotype than forwards(0.31 vs 0.28, respectively) and a lower proportion of the XX genotype (0.14 vs 0.19, respectively), also Heffernan et al., 2016 posits that the association of *ACTN3* R577X with playing position in elite rugby athletes suggests that inherited fatigue resistance is more prevalent in forwards, while inherited sprinting ability is more prevalent in backs. As for the *BDRKB2* gene, the difference is manifested in the predominance of the −9 allele referenced with vasodilation and cardiovascular performance, present more in backs.Similarly, in the *AGT* and IL6 genes, backs present a greater presence when genotypes such as CT and GG are related respectively, maintaining a genotypic predominance associated with muscle strength, reactivity and muscle recovery processes in the case of *IL6* (GG).

Among the characteristics to be highlighted are the low appearance of genotype XX (0.1) of the *ACTN3* gene, genotype II (0.02) of the *ACE* gene and genotype TT (0) of the *AGT* gene, which emphasizes a predominance of genotypes related to muscular strength. It is also appreciable the greater occurrence of athletes with production of ALPHA ACTININ 3 (*ACTN3* gene) either in homozygous form RR (0.574) or heterozygous RX (0.34) showing an artificial selection to the production of this protein, which is reported in studies linked to the Rugby Gen project (Bell et al., 2012, 2015; Heffernan et al., 2016) as a promoter of high performances in strength activities and protection to muscle damage in power efforts such as sprints, tackles, jumps and changes of direction.

## CONCLUSIONS

In conclusion, it is observed in the polymorphisms analyzed that their genotypic and allelic frequencies are associated according to the available literature with strength and power sports, and since rugby is a sport characterized as a sport of strength and speed, it is positively related as in the referenced studies. Also significant differences at genotypic and allelic level between the positions of backs and forwards game, although these differences are greater than those reported at least for ACE and ACTN3 genes, this may be due to the difference in levels of play, which in the studies related elite athletes and our study sub-elite athlete. in addition to a significant difference between the allelic and genotypic structure between athletes and control population that allows differentiating and establishing the athlete’s status.

At the same time, corroborating in a local study the association of different polymorphisms related to strength, power, aerobic capacity and muscle recovery in the international literature, especially when part of this is related to a genetic characterization project in rugby, such as the Rugby Gen project, allows us to use this information to specifically characterize a sport such as rugby in terms of its training processes and why not for selection processes and sports orientation.

## BIBLIOGRAPHY

Ahmetov, I. I., Emiliya, E., Egorova, S., Leysan, C., & Mustafina, J. (2013). the PPARA Gene Polymorphism in Team Sports Athletes. Central European Journal of Sport Sciences and Medicine, 1(1), 19–24.

Alves, C. R., Alves, G. B., Pereira, A. C., Trombetta, I. C., Dias, R. G., Mota, G. F. A., Fernandes, T., Krieger, J. E., Negrão, C. E., & Oliveira, E. M. (2013). Vascular reactivity and ACE activity response to exercise training are modulated by the +9/-9 bradykinin B2 receptor gene functional polymorphism. Physiological Genomics, 45(12), 487–492. https://doi.org/10.1152/physiolgenomics.00065.2012.

Andrade-Mayorga, O., Lavados-Romo, P., Valdebenito, C., Herrera, C. L., Carrasco, C., & Salazar, L. A. (2019). ACTN3 R577X Genetic Polymorphism in Chilean University Athletes. International Journal of Morphology, 37(4), 1493–1497. https://doi.org/10.4067/s0717-95022019000401493.

Bell, W., Colley, J. P., Evans, W. D., Darlington, S. E., & Cooper, S. M. (2012). ACTN3 genotypes of Rugby Union players: Distribution, power output and body composition. Annals of Human Biology, 39(1), 19–27. https://doi.org/10.3109/03014460.2011.632648.

Bell, W., Colley, J. P., Gwynne, J. R., Glazier, P., Evans, W. D., & Darlington, S. E. (2010). ACE ID genotype and leg power in Rugby Union players. Journal of Sports Medicine and Physical Fitness, 50(3), 350–355.

Bell, W., Colley, J. P., Evans, W. D., Darlington SE, Cooper, S. M., & Cobner, D. (2015). The ACTN3 Gene and Differences between Playing Positions in Bone Mineral Content, Fat Mass and Lean Tissue Mass in the Arms, Legs and Trunk of Rugby Union Football Players. Journal of Exercise, Sports & Orthopedics, 2(2), 1–7. https://doi.org/10.15226/2374-6904/2/2/00123. https://doi.org/10.15226/2374-6904/2/2/00123

Bell, W., Colley, J. P., Glazier, P., Evans, W. D., & Darlington, S. E. (2010). ACE ID genotype and leg power in Rugby Union players. September, 350–355.

Blair, M. R., Elsworthy, N., Rehrer, N. J., Button, C., & Gill, N. D. (2018). Physical and physiological demands of elite rugby union officials. International Journal of Sports Physiology and Performance, 13(9), 1199–1207. https://doi.org/10.1123/ijspp.2017-0849.

Brazier, J., Antrobus, M., Stebbings, G., Day, S., Callus, P., Erskine, R., Bennett, M., Kilduff, L., & Williams, A. (2018). Anthropometric and Physiological Characteristics of Elite Male Rugby Athletes. Journal OfStrength and Conditioning Research, 00(00), 1–12.

Brazier, J., Antrobus, M., Stebbings, G. K., Day, S. H., Heffernan, S. M., Cross, M. J., & Williams, A. G. (2019). Tendon and Ligament Injuries in Elite Rugby: The Potential Genetic Influence. Sports, 7(6), 138. https://doi.org/10.3390/sports7060138.

Cano, L. A., Piza, G., & Farfán, F. D. (2020). High intensity interval training in young rugby players from Argentina. Revista Internacional de Medicina y Ciencias de La Actividad Fisica y Del Deporte, 20(80), 505–512. https://doi.org/10.15366/rimcafd2020.80.002

CASAJUS, Juan. Determination of Work Rates in Rugby. Analysis of Records in First Division Teams. Period 1998-2001. G-SE Standard. 01/05/2003. g-se.com/a/82/.

Chiwaridzo, M., Oorschot, S., Dambi, J. M., Ferguson, G. D., Bonney, E., Mudawarima, T., Tadyanemhandu, C., & Smits-Engelsman, B. C. M. (2017). A systematic review investigating measurement properties of physiological tests in rugby. BMC Sports Science, Medicine and Rehabilitation, 9(1). https://doi.org/10.1186/s13102-017-0081-1. https://doi.org/10.1186/s13102-017-0081-1

Da Cruz-Ferreira, A. M., & Ribeiro, C. A. F. (2013). Anthropometric and physiological profile of portuguese rugby players - Part II: Comparison between athletes with different competitive levels. Revista Brasileira de Medicina Do Esporte, 19(1), 52–55. https://doi.org/10.1590/S1517-86922013000100011. https://doi.org/10.1590/S1517-86922013000100011.

Grant, D., David, P., & Sue, H. (2003). Applied Physiology and Game Analysis of Rugby Union. Sports Medicine, 33(13), 973–991. https://link-springer-com.chain.kent.ac.uk/content/pdf/10.2165%2F00007256-200333130-00003.pdf

Heffernan, S. M., Kilduff, L. P., Erskine, R. M., Day, S. H., McPhee, J. S., McMahon, G. E., Stebbings, G. K., Neale, J. P. H., Lockey, S. J., Ribbans, W. J., Cook, C. J., Vance, B., Raleigh, S. M., Roberts, C., Bennett, M. A., Wang, G., Collins, M., Pitsiladis, Y. P., & Williams, A. G. (2016). Association of ACTN3 R577X but not ACE I/D gene variants with elite rugby union player status and playing position. Physiological Genomics, 48(3), 196–201. https://doi.org/10.1152/physiolgenomics.00107.2015. https://doi.org/10.1152/physiolgenomics.00107.2015

Heffernan, S. M., Stebbings, G. K., Kilduff, L. P., Erskine, R. M., Day, S. H., Morse, C. I., McPhee, J. S., Cook, C. J., Vance, B., Ribbans, W. J., Raleigh, S. M., Roberts, C., Bennett, M. A., Wang, G., Collins, M., Pitsiladis, Y. P., & Williams, A. G. (2017). Fat mass and obesity associated (FTO) gene influences skeletal muscle phenotypes in non-resistance trained males and elite rugby playing position. BMC Genetics, 18(1). https://doi.org/10.1186/s12863-017-0470-1. https://doi.org/10.1186/s12863-017-0470-1

Heffernan, S. M., Kilduff, L. P., Day, S. H., Pitsiladis, Y. P., & Williams, A. G. (2015). Genomics in rugby union: A review and future prospects. European Journal of Sport Science, 15(6), 460–468. https://doi.org/10.1080/17461391.2015.1023222.

Heffernan, S. M., Kilduff, L. P., Erskine, R. M., Day, S. H., Stebbings, G. K., Cook, C. J., Raleigh, S. M., Bennett, M. A., Wang, G., Collins, M., Pitsiladis, Y. P., & Williams, A. G. (2017). COL5A1 gene variants previously associated with reduced soft tissue injury risk are associated with elite athlete status in rugby. BMC Genomics, 18(November). https://doi.org/10.1186/s12864-017-4187-3. https://doi.org/10.1186/s12864-017-4187-3

Johnson, W., Alderson, J., & Haff, G. (2017). Talent identification in elite rugby union: A theoretical update to an existing predictor algorithm. Journal of Australian Strength and Conditioning, 25(3), 33–41.

Lopes, A. L., Sant’Ana, R. T., Baroni, B. M., Cunha, G. dos S., Radaelli, R., Oliveira, Á. R. de, & Castro, F. de S. (2011). Anthropometric and physiological profile of Brazilian “rugby” athletes.” Revista Brasileira de Educação Física e Esporte, 25(3), 387–395. https://doi.org/10.1590/s1807-55092011000300004.

Massidda, M., Calò, C. M., Cieszczyk, P., Kikuchi, N., Ahmetov, I. I., & Williams, A. G. (2019). Genetics of team sports. In Sports, Exercise, and Nutritional Genomics: Current Status and Future Directions. https://doi.org/10.1016/B978-0-12-816193-7.00005-1.

McCormack, S., Jones, B., & Till, K. (2020). Training Practices of Academy Rugby League and their Alignment to Physical Qualities Deemed Important for Current and Future Performance. International Journal of Sports Science and Coaching, 15(4), 512–525. https://doi.org/10.1177/1747954120924905

Pickering, C., Kiely, J., Grgic, J., Lucia, A., & Del Coso, J. (2019). Can genetic testing identify talent for sport? Genes, 10(12). https://doi.org/10.3390/genes10120972.

Pimenta, E. M., Coelho, D. B., Cruz, I. R., Morandi, R. F., Veneroso, C. E., De Azambuja Pussieldi, G., Carvalho, M. R. S., Silami-Garcia, E., & De Paz Fernández, J. A. (2012). The ACTN3 genotype in soccer players in response to acute eccentric training. European Journal of Applied Physiology, 112(4), 1495–1503. https://doi.org/10.1007/s00421-011-2109-7.

Reilly, Thomas (1997). The physiology of rugby. Centre for Sport and Exercise Sciences, School of Human Sciences, John Moores University, Liverpool, England.

Sarzynski, M. A., Loos, R. J. F., Lucia, A., Pérusse, L., Roth, S. M., Wolfarth, B., Rankinen, T., & Bouchard, C. (2016). Advances in Exercise, Fitness, and Performance Genomics in 2015. Medicine and Science in Sports and Exercise, 48(10), 1906–1916. https://doi.org/10.1249/MSS.0000000000000982.

Suárez-Moreno Arrones, L.J.; Núñez, F.J. (2011). Physiological and anthropometric characteristics of elite rugby players in Spain and relative power out as predictor of performance in sprint and RSA. Journal of Sport and Health Research. 3(3):191–202

Szalata, M., Słomski, R., Balkó, Š., & Balkó, I. V. A. (2019). Advances in athlete genomics in 2019. Trends in Sport Sciences, 26(2), 55–61. https://doi.org/10.23829/TSS.2019.26.2-3. https://doi.org/10.23829/TSS.2019.26.2-3

Rett, K. (1990). Metabolic Effects of Kinins: Historical and Recent Developments. (p. 3). Journal cardiovascula Pharmacology Vol 15 1990.

Yamin, C., Duarte, J. A. R., Oliveira, J. M. F., Amir, O., Sagiv, M., Eynon, N., Sagiv, M., & Amir, R. E. (2008). IL6 (−174) and TNFA (−308) promoter polymorphisms are associated with systemic creatine kinase response to eccentric exercise. European Journal of Applied Physiology, 104(3), 579–586. https://doi.org/10.1007/s00421-008-0728-4

